# Application of Atlas of Cancer Signalling Network in pre-clinical studies

**DOI:** 10.1101/234823

**Authors:** L. Cristobal Monraz Gomez, Maria Kondratova, Jean-Marie Ravel, Emmanuel Barillot, Andrei Zinovyev, Inna Kuperstein

## Abstract

Initiation and progression of cancer involve multiple molecular mechanisms. The knowledge on these mechanisms is expanding and should be converted into guidelines for tackling the disease. We discuss here formalization of biological knowledge into a comprehensive resource Atlas of Cancer Signalling Network (ACSN) and Google Maps-based tool NaviCell that supports map navigation. The application of maps for omics data visualisation in the context of signalling maps is possible using NaviCell Web Service module and NaviCom tool for generation of network-based molecular portraits of cancer using multi-level omics data. We review how these resources and tools are applied for cancer pre-clinical studies among others for rationalizing synergistic effect of drugs and designing complex disease stage-specific druggable interventions following structural analysis of the maps together with omics data. Modules and maps of ACSN as signatures of biological functions, can help in cancer data analysis and interpretation. In addition, they can also be used to find association between perturbations in particular molecular mechanisms to the risk of a specific cancer type development. These approaches and beyond help to study interplay between molecular mechanisms of cancer, deciphering how gene interactions govern hallmarks of cancer in specific context. We discuss a perspective to develop a flexible methodology and a pipeline to enable systematic omics data analysis in the context of signalling network maps, for stratifying patients and suggesting interventions points and drug repositioning in cancer and other human diseases.

## Introduction

According to the current understanding, signalling pathways create a complex network with forward and backward regulatory loops, and many redundant pathways. The basic assumption is that during a pathological transformation cells do not create new signalling mechanisms, but rather hijack the existing molecular programs. This not only affects intracellular functions, but also the interactions between different cell types, leading to a new, yet pathological status of the system. There is a certain combination of molecular characteristics dictating specific cell signalling states enduring the pathological disease status. Identifying and manipulating the key molecular players controlling these cell signalling states, and shifting the pathological status toward the desired healthy phenotype are the major challenges for molecular biology of human diseases in general and in cancer, in particular [1][2][3].

To enable cancer data analysis in the context of molecular mechanisms, the information on these mechanisms should be systematically and adequately represented. The knowledge about molecular signalling mechanisms in cells is spread in thousands of publications, mostly in a human-readable form, hence precluding the application of bioinformatics and systems biology methods and algorithms. There is a need of formalized compilation of knowledge in a computer-readable form. The current solution is to represent the relationships between cellular molecules in a form of pathway diagrams, found in various pathway databases [4]. As the amount of information about biological mechanisms increases steadily, a different approach to organize and structure these data is essential. The aim is to create a more global picture of cell signalling with sufficient granularity for representation of molecular details, capturing cross-talks and feedback loops between molecular circuits. For this purpose, comprehensive signalling network maps covering multiple cellular processes simultaneously are more suitable than disconnected pathway diagrams. The approaches to rationally represent cell signalling in cancer, at the example of Atlas of Cancer Signalling Network (ACSN), are described in the review.

Visualisation and analysis of omics data in the context of signalling networks facilitates data interpretation and retrieval of deregulated mechanisms. Furthermore, data analysis in the context of signalling networks can help to detect patterns in the data projected onto the molecular mechanisms represented in the signalling maps, verifying enriched functional modules (‘hot’ deregulated areas), key players, and ‘bottleneck’ points [3] [5]. Correlating status of those network variables with the phenotype, as drug resistance or patient survival, followed by clustering methods, allows stratifying patients according to their integrated network-based molecular portraits and to design appropriate therapeutic intervention schemes [6].

As a productive idea for developing intervention schemes, synthetic lethality (SL) provides a conceptual framework for the development of cancer-specific drugs. The classical paradigm defines synthetic lethal interactions as a phenomena where combinations of two gene deletions significantly affects cell viability, whereas single deletion of each one of those genes does not [7]. The idea of SL treatment approach is to take an advantage of the specificities in tumour cells which bearing abnormal function of one of the genes from the synthetic lethal pair. Targeting synthetic lethal partner allows then selective killing of tumour cells, and avoiding or limiting side effects on normal cells [8], and it also provides the clinician with a biomarker for selecting patients that expected to respond to the treatment. Giving the complexity of signalling mechanisms simultaneously involved in cancer, the synthetic lethality pairs paradigm should be extended to the synthetic lethal sets or combinations paradigm [9] [10]. The computational approaches that allow *in silico* testing of multiple synthetic interactions combinations, considering large comprehensive signalling networks and cancer omics data are discussed in this review.

### Atlas of Cancer Signalling Network: geographical map of molecular mechanisms

Deregulation of molecular mechanisms leading to cancer pertain to various processes such as cell cycle, cell death, DNA repair and DNA replication, cell motility and adhesion, cell survival mechanisms, immune processes, angiogenesis, tumour microenvironment and others. Most of them are collectively or sequentially involved in tumour formation and modified as the tumour evolves. It is assumed that in pathological situations the normal cell signalling network is altered by deregulated coordination between pathways or disruption of existing molecular pathways, rather than by creating completely new signalling pathways and molecular interactions. The most common abnormalities in pathological situations are perturbations at the gene expression level, protein abundance or protein post-translational modifications, irregular ‘firing’ or silencing of particular signals, wrong sub-cellular localization of particular molecules and so on. Such quantitative rather than qualitative network changes compared with normal cell signalling could be studied in the context of comprehensive signalling networks by analysing experimental data obtained from tumour samples, patient-derived xenografts, cancer-related cell lines or animal models. This approach helps to understand the interplay between molecular mechanisms in cancer, deciphering how gene interactions govern hallmarks of cancer [11] in specific settings.

Despite the existence of a large variety of pathway databases and resources [4], only few of them depict processes specifically implicated in cancer and none of those resources depict the processes with enough granularity. In addition, pathway browsing interfaces play more and more important role in cancer research, but require further improvements. Therefore, we constructed a resource Atlas of Cancer of Signalling Networks (ACSN, http://acsn.curie.fr) that aims to formalize knowledge on cancer-related processes in a form of a comprehensive signalling network map for data interpretation in basic research and clinical studies [12].

The construction and update of ACSN involve manual mining of molecular biology literature along with the participation of experts in those fields. ACSN differs from other databases because it contains a deep comprehensive description of cancer-related mechanisms retrieved from the most recent literature, following the hallmarks of cancer (Figure 1). Cell signalling mechanisms are depicted using the CellDesigner tool [13] at the level of biochemical interactions, assembling a large network of 4826 reactions covering 2371 proteins and based on approximately 3000 references (Table 1).

**Figure 1.**
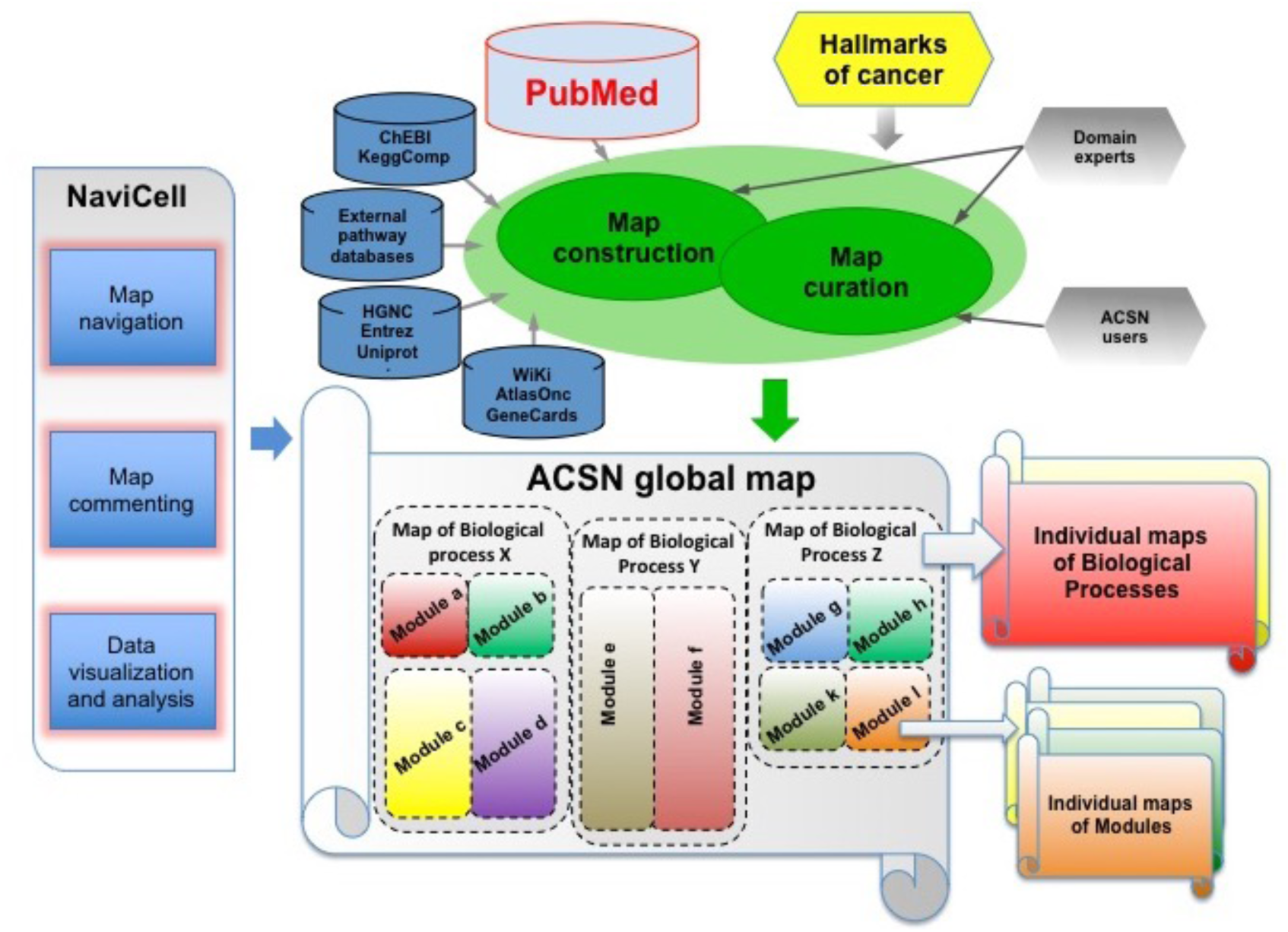
*Structure of Atlas of Cancer Signalling Network resource*. *The scheme demonstrates the concept of ACSN construction starting from the cancer hallmarks: collecting information about molecular mechanisms underlying those hallmarks from scientific publications and manually depicting them in the global map of ACSN and further supporting by consulting the information from the external pathway databases. ACSN is hierarchically organized into three levels: the seamless global map divided into the interconnected biological process maps that are further decomposed into interconnected module maps. ACSN can be exploited through web-based NaviCell interface allowing map navigation using Google Maps engine, map commenting via associated blog system and user omics data visualisation and analysis (Adapted from* [12]).

**Table 1:** *Content of ACSN (Adapted from [13])*

| Feature | Content |
| --- | --- |
| Maps of biological processes | 5 |
| Functional modules | 52 |
| Chemical species | 5975 |
| Reactions | 4826 |
| Proteins | 2371 |
| Metabolites | 595 |
| Genes | 159 |
| References | 2919 |

Currently ACSN contains representations of molecular mechanisms that are frequently deregulated in cancer, such as cell cycle, DNA repair, cell death, cell survival, and epithelial to mesenchymal transition (EMT). Cell signalling mechanisms are depicted on the maps in great detail, creating together a seamless map of molecular interactions, presented as a global ‘geographic-like’ molecular map (Figure 1A). ACSN has a hierarchical structure, composed of interconnected maps of biological process implicated in cancer. Each map is further divided into functional modules, corresponding mainly to canonical signalling pathways (Figure 1, 2C).

**Figure 2.**
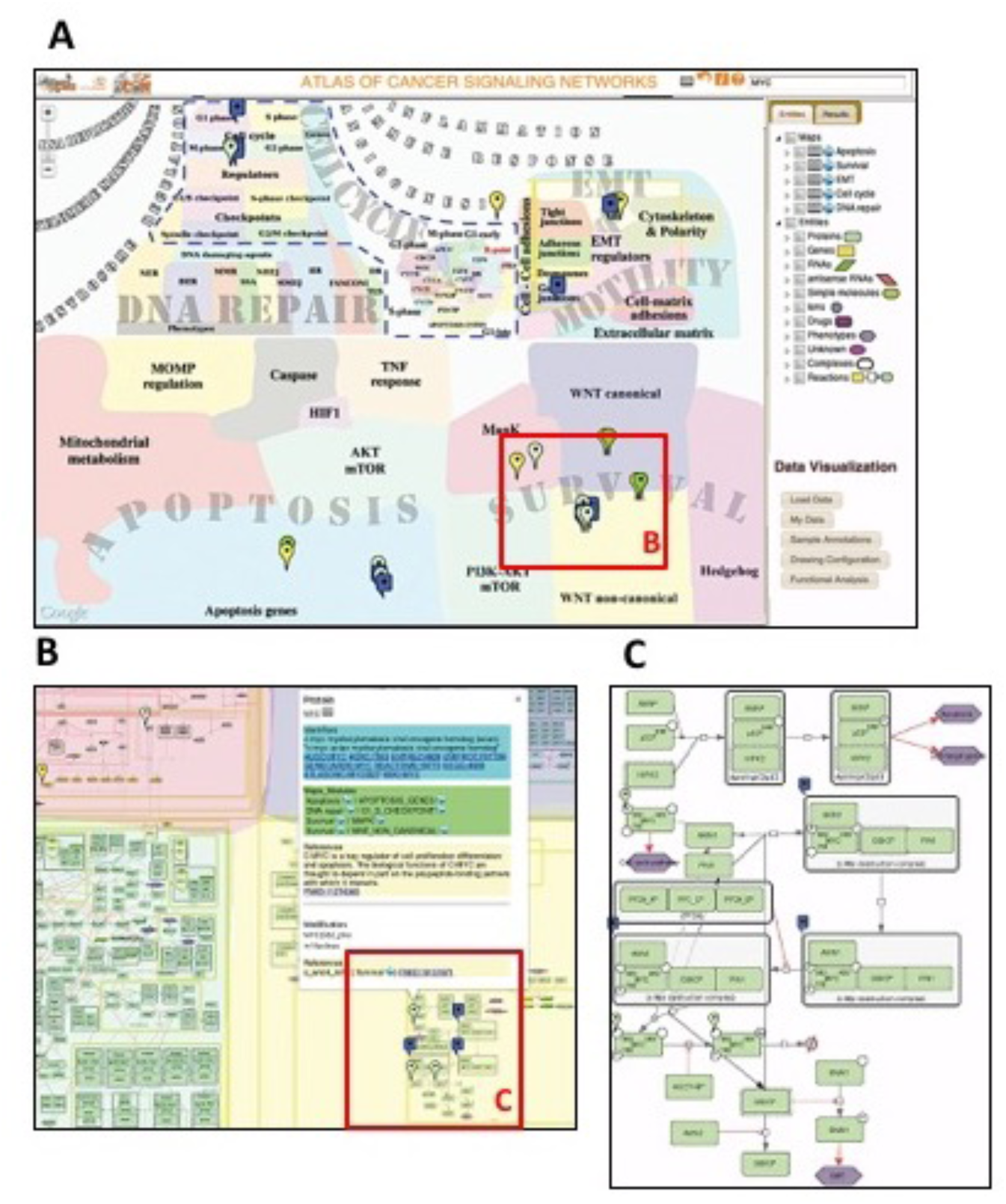
*Browsing interface of Atlas of Cancer Signalling Network*. *(A). ACSN interface with selection panel and data visualisation menu. Querying ACSN is possible via the search window or by checking on the entity in the list of entities. Distribution of frequently mutated oncogene MYC across molecular mechanisms on the ACSN maps is indicated; (B). Google Maps-like features of NaviCell for visualisation and annotation of map entities (markers, callout with links to external databases, citations and ACSN maps of and functional modules where MYC protein is found); (C). Zoom in on Wnt non-canonical module of cell survival map to observe signalling processes where MYC protein is involved (Adapted from* [12]).

The navigation interface include features such as scrolling, zooming, markers and callouts using Google Maps technology adapted by NaviCell [14] (Figure 3), web-based platform supporting ACSN and similar efforts in CellDesigner format [15,16] or other formats [17]. The semantic zooming in NaviCell (http://navicell.curie.fr), provides several view levels, achieved by gradual exclusion of details and abstraction of information upon zooming out (Figure 2B).

**Figure 3.**
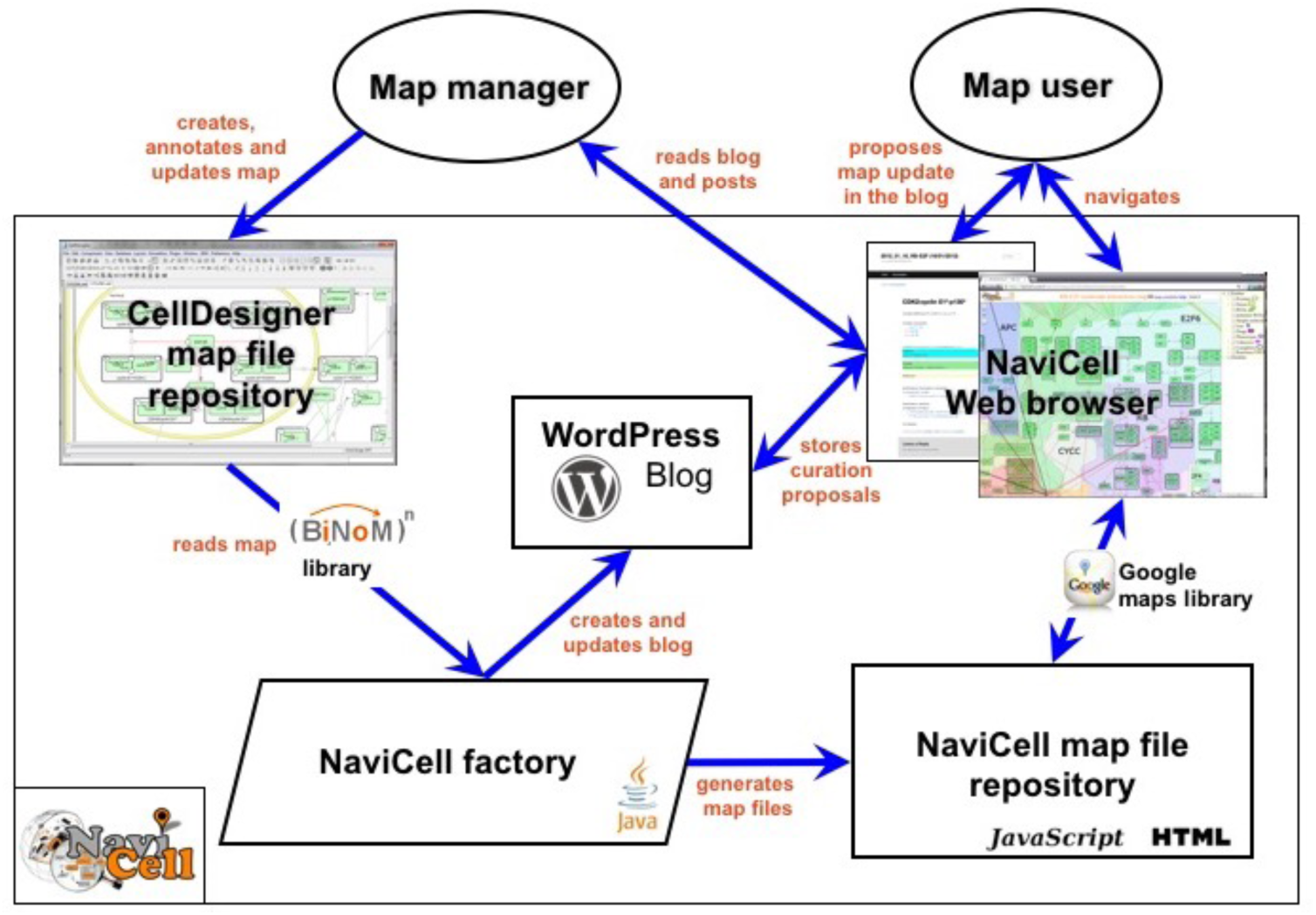
*General architecture of NaviCell environment (Adapted from* [18]).

ACSN is a unique resource of cancer signalling, with an enormous amount of information embedded and organized. Together with NaviCell, it is optimized for integration and visualisation of cancer molecular profiles generated by high-throughput techniques, drug screening data or synthetic interactions studies. The integration and analysis of these data in the context of ACSN may help to better understand the biological significance of results, guiding scientific hypotheses and suggesting potential intervention points for cancer patients. In addition, since ACSN covers major cell signalling processes, the resource and associated methods for data analysis using ACSN are suitable for applications in many biological fields and for studying various human diseases.

The atlas is currently being extended with additional maps depicting molecular mechanisms of DNA replication, telomere maintenance, angiogenesis, immune response and others that will be integrated in future releases. The atlas will not only cover intracellular processes, but also cross talk of cancer cell with the components of tumour microenvironment. An additional level of complexity will be added to the atlas in near future, representing different types of cells surrounding tumour, and their interplay, to enable modelling of complex phenotypes.

### Molecular portraits of cancer: data visualisation and analysis using Atlas of Cancer Signalling Network

#### Data visualisation in NaviCell Web Service environment

The data integration into ACSN is possible using NaviCell Web Service, an user-friendly environment embedded into the NaviCell tool [18]. It allows uploading several types of “omics” data; e.g. expression data for mRNA, microRNA, proteins, mutation profiles, copy-number data, and visualize them simultaneously in the context of molecular interaction maps (Figure 4).

**Figure 4.**
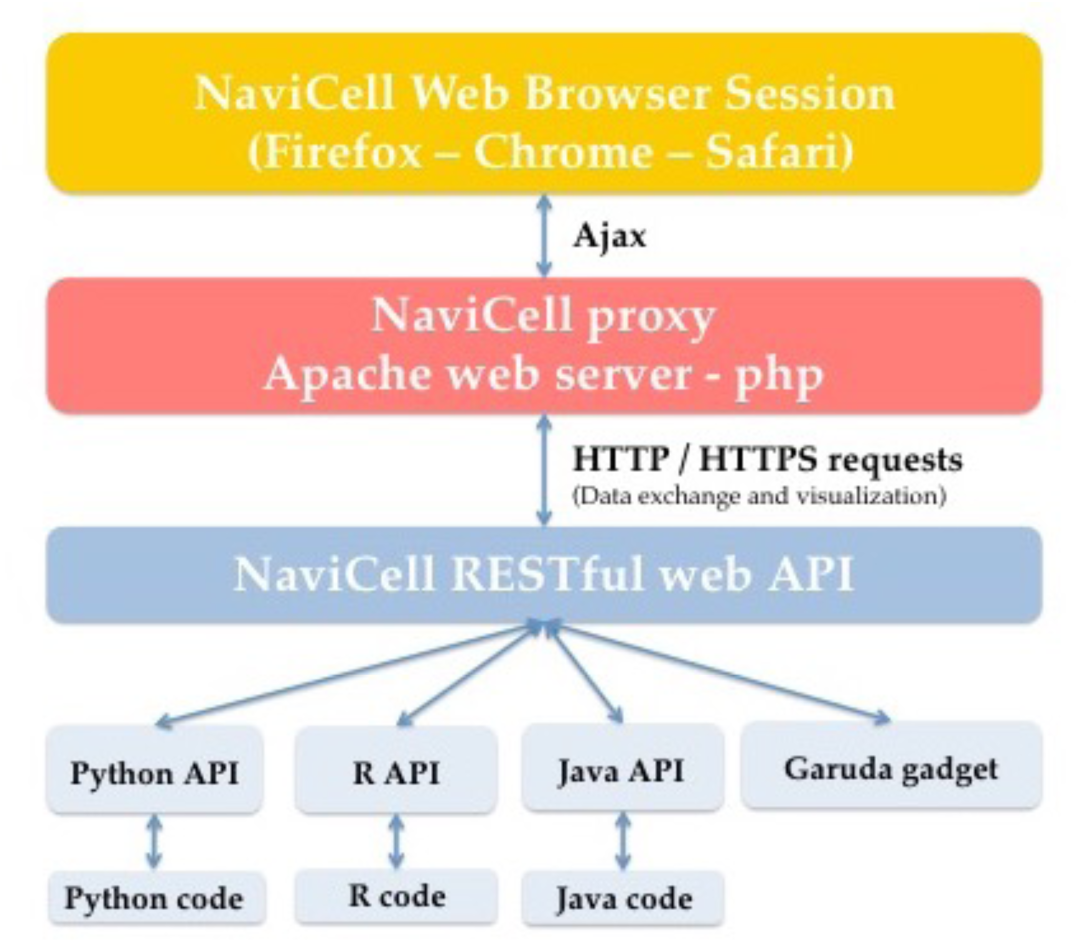
*General architecture of NaviCell Web service server*. *Client software (light blue layer) communicates with the server (red layer) through standard HTTP requests using the standard JSON format to encode data (RESTful web service, dark blue layer). A session (with a unique ID) is established between the server and the browser (yellow layer) through Ajax communication channel to visualize the results of the commands send by the software client (Adapted from* [18]).

Depending on the nature of the data, different types of visualisation modes can be required to achieve the informative picture maps, bar plots, glyphs and map staining (Table 2). The data can be visualized at different zoom levels. Sample annotation files uploaded with the data can serve to define groups of samples.

**Table 2.**
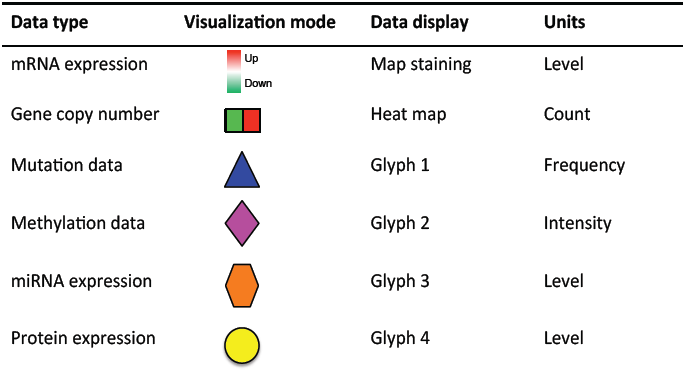
*Data display modes in NaviCell and NaviCom (Adapted from* [19]).

| Data type | Visualization mode | Data display | Units |
| --- | --- | --- | --- |
| mRNA expression | Up<br>Down | Map staining | Level |
| Gene copy number |  | Heat map | Count |
| Mutation data |  | Glyph 1 | Frequency |
| Methylation data |  | Glyph 2 | Intensity |
| miRNA expression |  | Glyph 3 | Level |
| Protein expression |  | Glyph 4 | Level |

A novel mode of data visualisation for continuous data (e.g. expression) provided by NaviCell Web Service is the ‘map staining’. In this technique, the values mapped to individual molecular entities or group of entities (e.g. score of functional module activities, see below), results in a colourful background of the network map that represents the data distribution pattern [18]. All those approaches for data integration into the signalling maps described above allow to rationalize the information embedded into the data: compare samples or group of samples; find typical patterns of data distribution across the molecular mechanisms depicted on the maps; grasp deregulated 'hot area' on the maps and major involved players and draw hypothesis as to which mechanisms to concentrate the work in the studied samples. These signalling network-based molecular signatures of samples thus help to stratify patients (Figure 5).

**Figure 5.**
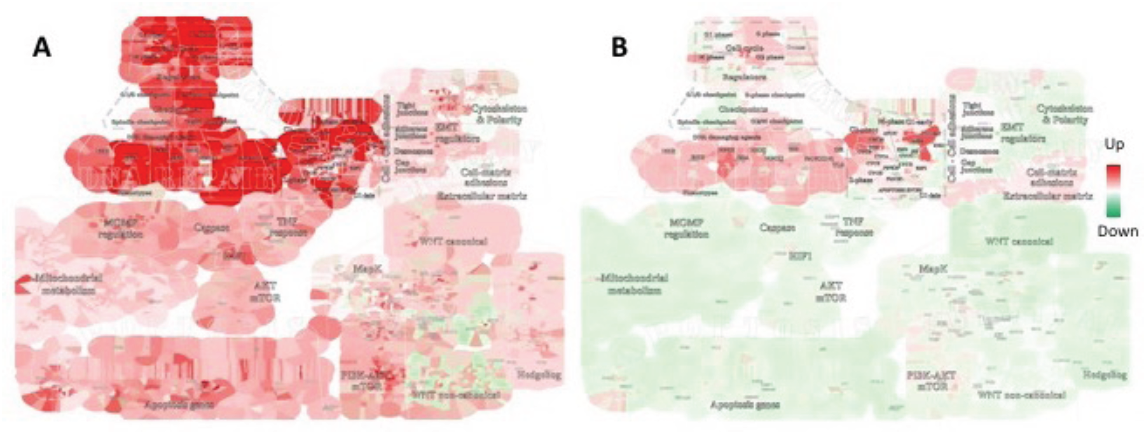
*Breast cancer gene expression data integration and analysis using NaviCell.* *The mRNA expression data from TCGA collection has been used for evaluation of functional modules activities and ACSN coloring as ‘map staining’ for (A) Basal-like and (B) Luminal A breast cancer types. The two breast cancer subtypes are characterized by different patterns of module activities. (Adapted from* [12]).

Various omics data are available on the public and local databases [20]. However, there are no tools supporting the import of big datasets from these databases and display on signalling network maps in an efficient manner and with optimized visualisation settings. To answer to this demand, NaviCom has been developed, a python package and web interface for automatic simultaneous display of multi-level data in the context of signalling network map [19]. NaviCom (http://navicom.curie.fr) is bridging between the cBioPortal database and the NaviCell interactive tool, for data visualisation NaviCom is empowered by a cBioFetchR R package to import high-throughput data sets from cBioPortal to NaviCell and Navicom Python module, permitting automatized simultaneous visualisation of multilevel omics data on the interactive signalling network maps using NaviCell environment (Figure 6A). NaviCom proposes several standardized modes of data display on signalling networks maps to address specific biological questions (Table 2).

**Figure 6.**
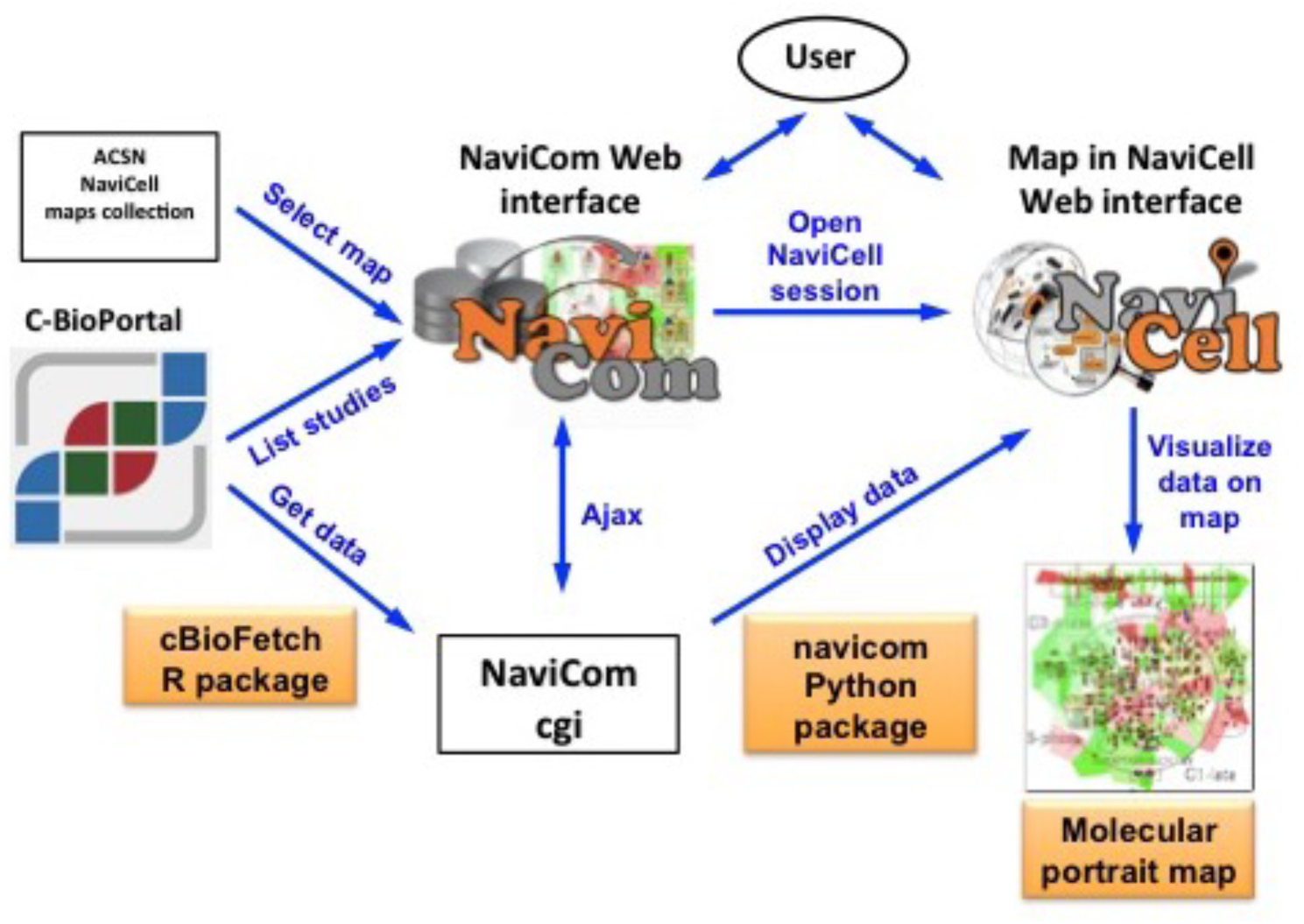
*General architecture of NaviCom environment*. *The NaviCom interface provides the user with an updated list of studies from cBioPortal and links to ACSN and NaviCell maps collections. When visualisation is launched, NaviCom starts a new NaviCell session and calls a cgi on the server. The cgi downloads cBioPortal data to the NaviCell session and displays them to generate the molecular portrait selected by the user. (Adapted from* [19]).

This tool enables generation of complex molecular portraits from multiple omics datasets from cBioPortal. In near future the NaviCom platform will be extended and will provide access to any type of omics data from a wide range of databases (TCGA, ICGC, HGMB, METABRIC, CCLE). In addition, to allow broader description of molecular mechanisms implicated in the studied sample, signalling networks available in databases as KEGG [21], Reactome [22] and others, will be also integrated and used for high-throughput data analysis via NaviCom platform.

#### Data analysis using ACSN

To identify enriched modules of a molecular map from a given dataset, several tools are currently available. Gene Set Enrichment Analysis (GSEA) (http://software.broadinstitute.org/gsea/index.jsp) is a computational method aimed to find over-represented modules in a ranked gene list, using a weighted Kolmogorov-Smirnov test [23]. ACSNMineR (https://github.com/sysbio-curie/ACSNMineR) is an R package that incorporates ACSN information to calculate enriched or depleted modules by means of a Fisher exact test or a hypergeometric test [24]. The Representation and Quantification of Module Activity (ROMA) method, implemented in Java and R (https://github.com/sysbio-curie/Roma, https://github.com/sysbio-curie/rRoma, https://github.com/sysbio-curie/rRoma), designed for fast and robust computation of the activity of gene sets (or modules) with coordinated expression. ROMA is using the first principal component of a PCA analysis to summarize the coexpression of a group of genes in the gene set. ROMA is also providing additional functionalities: (i) calculation of the individual gene contribution to the module activity level and determination of the genes that are contributing the most to the first pricipal component, (ii) alternative techniques to compute the first principal component, i.e. weighted and centered methods, (iii) estimation of the statistical significance of the proportion of variance explained by the first principal component, as well as the spectral gap between the variance explained by the first and second component (representing the homogeneity for the gene set) [25]. The module activity scores calculated by these methods, can be visualized in the context of ACSN using the map staining technique as described above (Figure 5). Such visualization is automated in ACSNMiner and ROMA.

ACSN was also used, as a source of module definitions for benchmarking DeDaL tool. DeDaL provides a possibility to create data-driven and structure-driven network layouts, which are more insightful for grasping correlation patterns in multivariate data on top of networks [26]. ACSN module definitions were applied for testing a method for inferring hidden causal relations between pathway members using reduced Google matrix of directed biological networks [27].

ACSN serves as a source of functional module definitions and PPI network in the data analysis projects, especially those dedicated to cancer data analysis. For example, gene lists from functional modules of DNA repair map were used to study homologous recombination deficiency in invasive breast carcinomas[28]. ACSN was used as a source of signatures for processes involved in cancer for classification of gene signatures and generation of InfoSigMap, an interactive online map showing the structure of compositional and functional redundancies between signatures from various sources [29].

### Exploiting the Atlas of Cancer Signalling Network in pre-clinical research

#### Explaining synergistic effect of combined treatment in breast cancer

DNA repair inhibitors are holding promises to improve cancer therapy but their application is limited by the compensatory activities of different repair pathways in cancer cells. For example, PARP inhibitors that act as synthetic lethal with BRCA deficiency, appear however less efficient in patients with active Homologous Recombination (HR) repair [30]. During treatment, some tumours escape through compensatory mutations that restore the HR activity or stimulate the activity of alternative repair pathways such as Non-Homologous End Joining (NHEJ) and Alternative Non-Homologous End Joining (Alt-NHEJ). A new class of DNA repair pathways inhibitors (Dbait or AsiDNA, the derivative of Dbait) have been recently developed, consisting of 32bp deoxyribonucleotides DNA double helix that mimics double strand breaks (DSB). It acts as an agonist of DNA damage signalling thereby inhibiting DNA repair enzyme recruitment at the damage site [31]. However, studies of Dbait effects on multiple types of cancer cell lines show occurrences of resistance in cancer type-independent manner.

Depending on the genetic background, different breast cancer tumours vary in their sensitivity to DNA repair inhibitors, as PARP inhibitors and Dbait. To understand molecular mechanisms underlining these differences, a combination of experimental and bioinformatics approaches was applied. Triple Negative Breast Cancer (TNBC) cell lines were studied for their sensitivity to the AsiDNA, the derivative of Dbait DNA repair inhibitior and Olaparib, the PARP inhibitor. Different TNBC cell lines show wide distribution of response/resistance to these drugs, despite that fact the cell lines are related to the same disease type. Integrative analysis of omics data from these cell lines covering mRNA expression, copy number variations and mutational profiles was performed, retrieving non-overlapping unique gene sets robustly correlated with resistance to each one of the drugs. Analysis of the omics data in the context of ACSN maps highlighted deregulated functional modules across ACSN, associated with resistance to each one of the drugs allowing to established drug resistance network-based molecular portraits. This analysis confirmed that different specific defects in DNA repair machinery are associated to AsiDNA (Figure7A) or Olaparib (Figure7B) resistance. Importantly, it showed involvement of different compensatory DNA repair mechanisms in cell lines resistant to AsiDNA comparing to cell lines resistant to Olaparib (Figure7D), suggesting a rational for combination of these two drugs. The authors confirmed synergistic therapeutic effect of the combined treatment with AsiDNA and PARP inhibitors in TNBC, while sparing healthy tissue (Figure 7C) [32].

**Figure 7.**
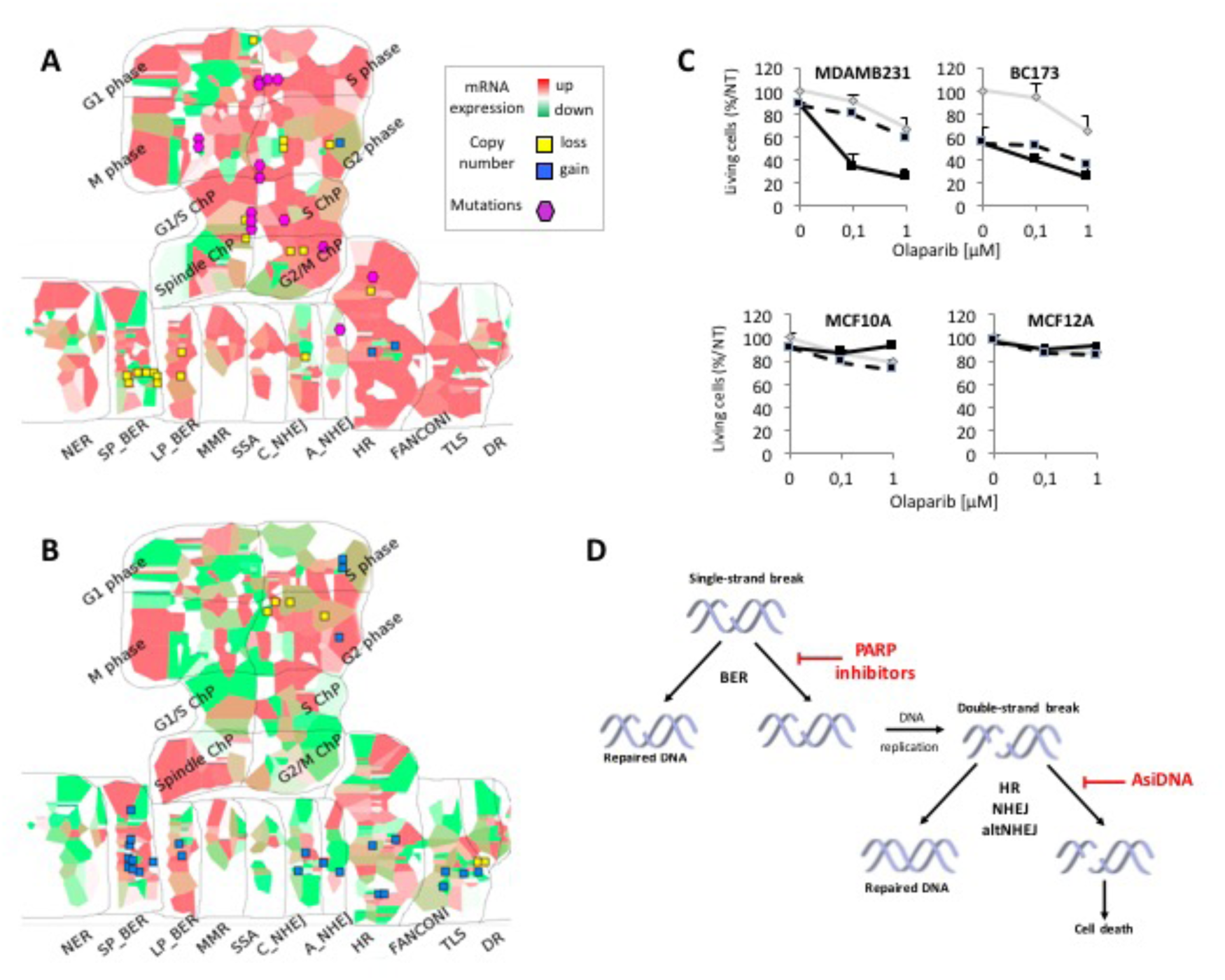
*Overcoming resistance of Triple Negative Breast Cancer cell lines to DNA repair inhibitors*. *Molecular portraits of TNBC cell lines resistant to (A) AsiDNA or (B) Olaparib, visualized on DNA repair map. (C). Cell survival to combination of AsiDNA and Olaparib, with DT01(black line), without DT01(grey line); dashed lines indicate calculated cell survival for additive effect of two drugs. (D) Schematic representation of inhibitory mechanisms of AsiDNA and Olaparib. Base Excision Repair (BER), Homologous Recombination (HR), Non-Homologous End Joining (NHEJ), Alternative Non-Homologous End Joining (Alt-NHEJ). (Adapted from* [32]).

#### Complex stage-specific interventions in MAPK pathway to disrupt proliferative signalling in bladder cancer

The idea of synthetic lethality (SL) treatment approach is to take an advantage of the specificities in tumour cells displaying abnormal expression or function of one gene from a synthetic lethal pair. Targeting synthetic lethal partner allows then to selective killing of tumour cells [33]. This approach is applied in BRCA2 mutated breast cancer cases using PARP inhibitors, however there is frequent escape from the treatment, requiring of a more complex solution. Treatment failure can be due to robustness of cell signalling network ensured by redundant mechanisms that provide the possibility to bypass the effect of drugs [34]. Therefore, the ways for identifying and blocking those active compensatory pathways should be found. One of the approaches is taking into account the signalling network structure to find the most optimal SL combinations of genes, probably more than a pair [10][35][36].

The computational strategy to suggest complex intervention sets has been developed and demonstrated using MAPK signalling network [37]. MAPK signalling network is coordinating between various processes implicated in cell survival and currently included into the Cell Survival map of ACSN. The strategy involves two steps: i) identification of tumour stage-specific active functional modules, i.e. sets of MAPK signalling network components that are transcriptionally deregulated in bladder cancer [38] compared to normal samples, and ii) computation of intervention sets of MAPK map components, whose disruption block all the proliferative paths fostered by the identified active functional modules in bladder cancer [39] (Figure 8A).

**Figure 8.**
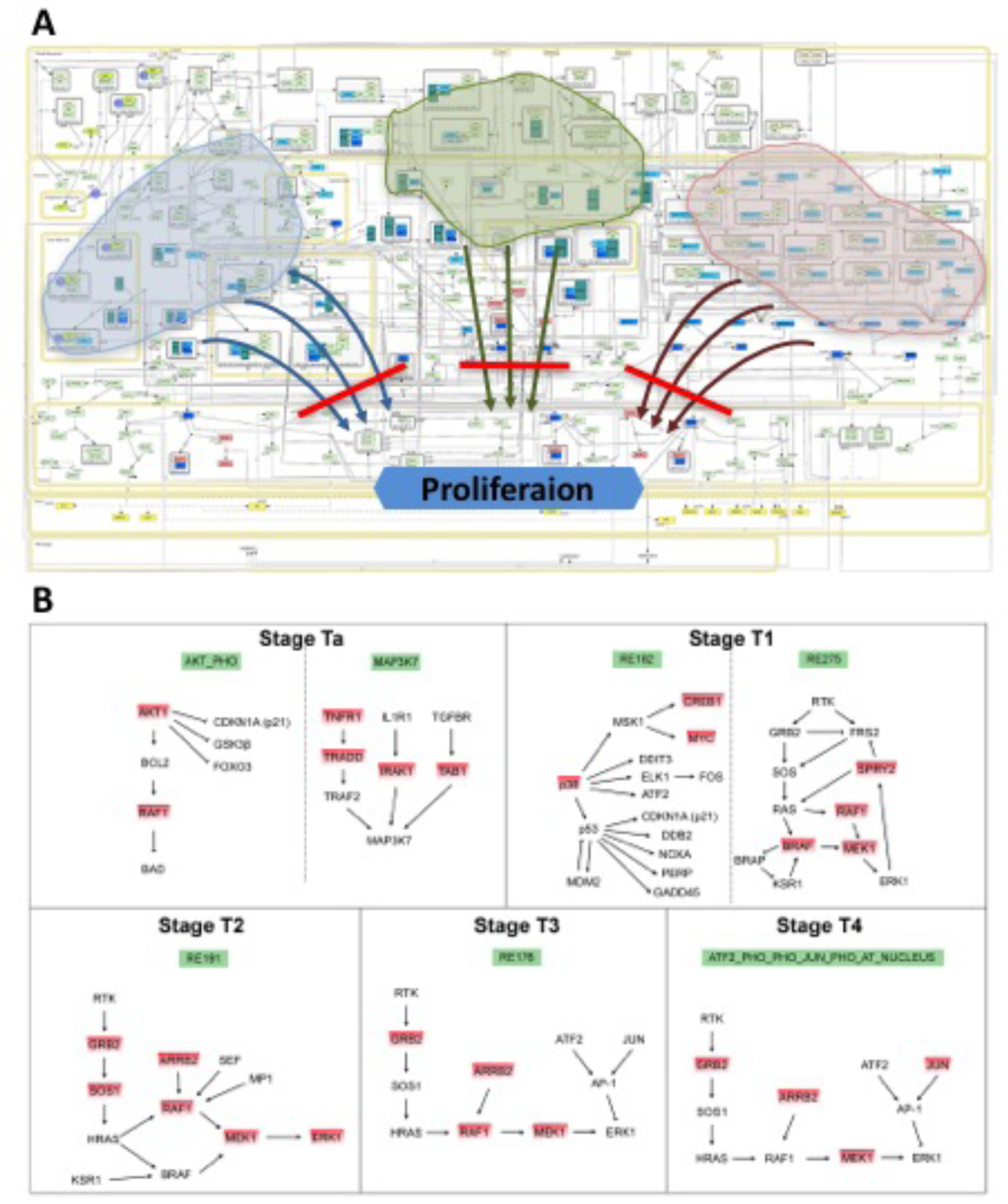
*Computational strategy for finding stage-specific interventions sets using detailed reaction network analysis and omics data*. *(A) Detailed MAPK network map is shown with schematically indicated activated modules (see text). Finding minimal hitting sets allows to cut all paths (schematically shown by arrows) from the activated modules to the proliferation phenotype. (B) Stage-specific activated modules detected in MAPK network using bladder cancer transcriptome data. Module enrichment score were computed by GSEA method. The most contributing leading-edge genes with highest differential expression level are highlighted by red. The optimal hitting set lists from these genes elements of MAPK network, which removal cuts all the paths from the corresponding activated modules to the proliferation phenotype were calculated using OCSANA algorithm and interventions sets for each stage of bladder cancer were suggested (see text). (Adapted from* [39]).

The procedure was applied to five different bladder tumour stages [38]. Differential gene expression levels were computed relative to healthy conditions, using expression data. Regions of the map having high density of differentially expressed genes/proteins were identified and scored by Gene Set Enrichment Analysis (GSEA) [23]. The highly scored regions were assumed likely to point to sources of proliferative signals in the tumour. If paths exist in the map from the components belonging to any strongly activated network region to the node “Proliferation”, then presumably the region contributes to the activation of cell division (Figure 8A). The removal of a set of proteins from the network might block all such proliferative paths. This intuition was formalized by the notion of the minimal cut sets, which were computed using the OCSANA algorithm in Cytoscape plugin BiNoM [40]. OCSANA algorithm computes the minimal cut sets by simultaneously prioritizing them with respect to the potential effect on the target network nodes while avoiding side effects on the parts of the network which function should be preserved.

In the less invasive Ta stage, two significantly activated functional modules were identified (Figure 8B; stages Ta). One contains AKT from the PI3K pathway, that has been shown to be deeply involved in bladder cancer [41], inhibitors targeting this protein have been recently developed [42]. As OCSANA intervention set the analysis suggests (Table 3), RAS de-phosphorylation should inhibit propagation of signal through this module to proliferation in Ta stage tumour. The second module contains MAP3K7 protein, an upstream activator of p38 and JNK, which is activated by three stimuli: TNFR1, IL1R1 and TGFβR. All these pathways are frequently up-regulated in Ta tumours. OCSANA results suggested that the de-phosphorylation of both p38 and MAP3K7 can block the proliferative effects of MAP3K7-dependent functional module. Interestingly, bladder cancer cell lines were shown to proliferate due to the joint activity of PI3K and p38 (unpublished data), especially when FGFR3 is active. In the dataset considered for the analysis, FGFR3 gene is strongly expressed in stage Ta, less in stage T1, whereas it has low expression in invasive bladder tumours. Strikingly, the best intervention for Ta tumours consists of the disruption of both p38 and a downstream target of PI3K (i.e. AKT).

**Table 3.**
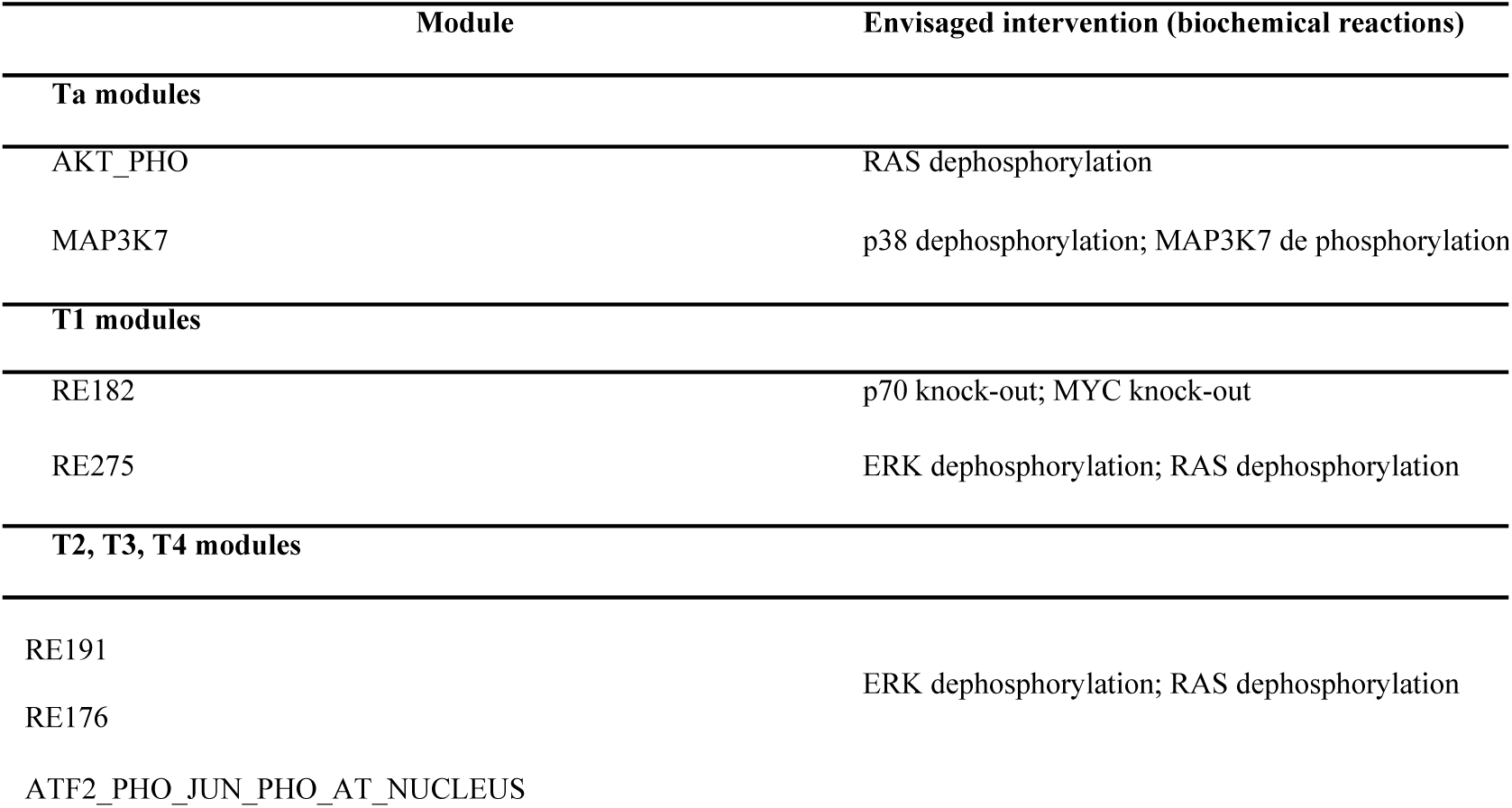
*Intervention sets for stage-specific activated modules in bladder cancer (Adapted from* [39]).

In the most invasive T2, T3 and T4 stages of bladder cancer there are three up-regulated functional modules, all characterised by high expression of RTK/ERK signalling components (Figure 8B; stages T2, T3 and T4). OCSANA results (Table 3) point to the de-phosphorylation of both ERK and RAS as best interventions to block proliferation, which makes it coherent the current developments of RAS- and ERK-inhibiting drugs for several cancer types, including bladder cancer[43].

The analysis suggested different interventions depending on tumour stage. In less invasive tumours p38 coupled with PI3K-dependent signalling could be targeted, whereas RAS/ERK pathway is likely more critical for more invasive stages. Similarly, network analysis using ACSN and OCSANA can be performed for individual patient tumour profiles, leading to personalized recommendations of treatment [44].

#### Finding metastasis inducers in colon cancer through network analysis

Evolution of invasion and metastasis, in particular in colon cancer, has been studied in experimental models, however, the mechanism that triggers the process is still not clear and the available mice models of colon cancer are far from being satisfactory [45][46]. With the aim to create an experimental mouse model of invasive colon cancer, one needs to address the question what are the major players and the driver mutations inducing invasion. One of early events of metastasis is assumed to be epithelial to mesenchymal transition (EMT) [47].

In order to identify the interplay between signalling pathways regulating EMT, a signalling network was manually created based on the information retrieved from around 200 publications. This signalling map is integrated into the EMT and cell motility comprehensive map of ACSN. Structural analysis and simplification of the EMT network that highlighted the following EMT network organisation principles, which is in agreement with current EMT understanding: (1) Five EMT transcription factors SNAIL, SLUG, TWIST, ZEB1 and ZEB2 that have partially overlapping sets of downstream target genes can activate the EMT-like program. (2) These key EMT transcription factors are under control of several upstream mechanisms: they are directly induced at the transcriptional level by the activated form of Notch, Notch Intracellular Domain (NICD), but are downregulated at the translational level by several miRNAs that are under transcriptional control of p53 family genes. (3) All five key EMT transcription factors should be activated ensuring simultaneous activation of EMT-like programme genes and downregulating miRNAs. In addition, the EMT key inducers also inhibit apoptosis and reduce proliferation. (4) The activity of Wnt pathway is stimulated by transcriptional activation of the gene coding for beta-catenin protein by NICD-induced TWIST or SNAI1. The Wnt pathway, in turn, can induce the expression of Notch pathway factors, creating a positive feedback loop. (5) Components of the Wnt and Notch pathways are negatively regulated by miRNAs induced by the p53 family (p53, p63 and p73). The balance between the effect of positive (Notch and Wnt) and negative (p53, p63 and p73 mediated by miRNAs) regulatory circuits on EMT inducers dictates the possibility of EMT phenotype [48][49][50][51]. Based on those features, the hub players were highlighted and network complexity reduction was performed using Cytoscape plugin BiNoM. The reduced network contained the core regulatory cascades of EMT, apoptosis and proliferation that were preserved through all levels of reduction [52]. This reduced network has been used for comparison between the wild type and all possible combinations of single and double mutants for achieving EMT phenotype. The computational analysis of the signalling network lead to the prediction that the simultaneous activation of NICD and loss of p53 can promote an EMT phenotype. Furthermore, EMT inducers may activate the Wnt pathway, possibly resulting in a positive feedback loop that will amplify Notch activation and maintain an EMT-like program (Figure 9).

**Figure 9.**
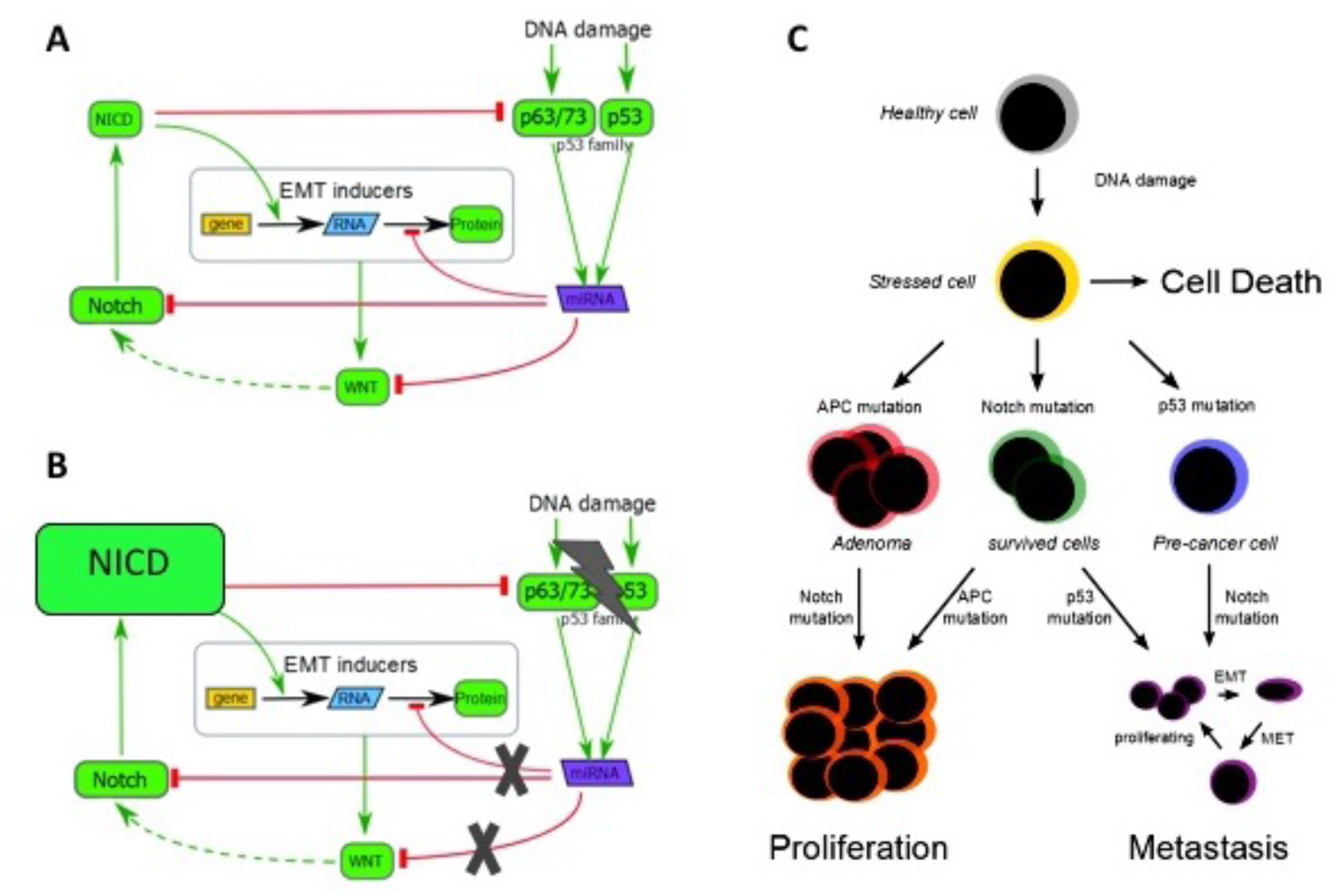
*Prediction of synthetic interaction combination to achieve EMT*. *Mechanistic model of EMT inducers regulation involving Notch (NICD), p53 and Wnt pathways in (A) normal and (B) double mutant with NICD overexpressed and p53 lost. (C) Scheme representing regulation of three major cell states in colon cancer (Cell death, Proliferation, Metastasis). (Adapted from* [53]).

To validate this hypothesis, a transgenic mouse model was generated, expressing a constitutively active Notch1 receptor in a p53-deleted background, specifically in the digestive epithelium. Importantly, green fluorescent protein (GFP) expression linked to the Notch1 receptor activation allows lineage tracing of epithelial tumour cells during cancer progression and invasion (Figure 10A). These mice develop digestive tumours with dissemination of EMT-like epithelial malignant cells to the lymph nodes, liver and peritoneum and generation of distant metastases (Figure 10B). Exploration of early EMT program inducers in in invasive human colon cancer samples confirmed that EMT markers are associated with modulation of Notch and p53 gene expression in similar manner as in the mice model (Figure 10C), supporting a synergy between these genes to permit EMT induction [53].

**Figure 10.**
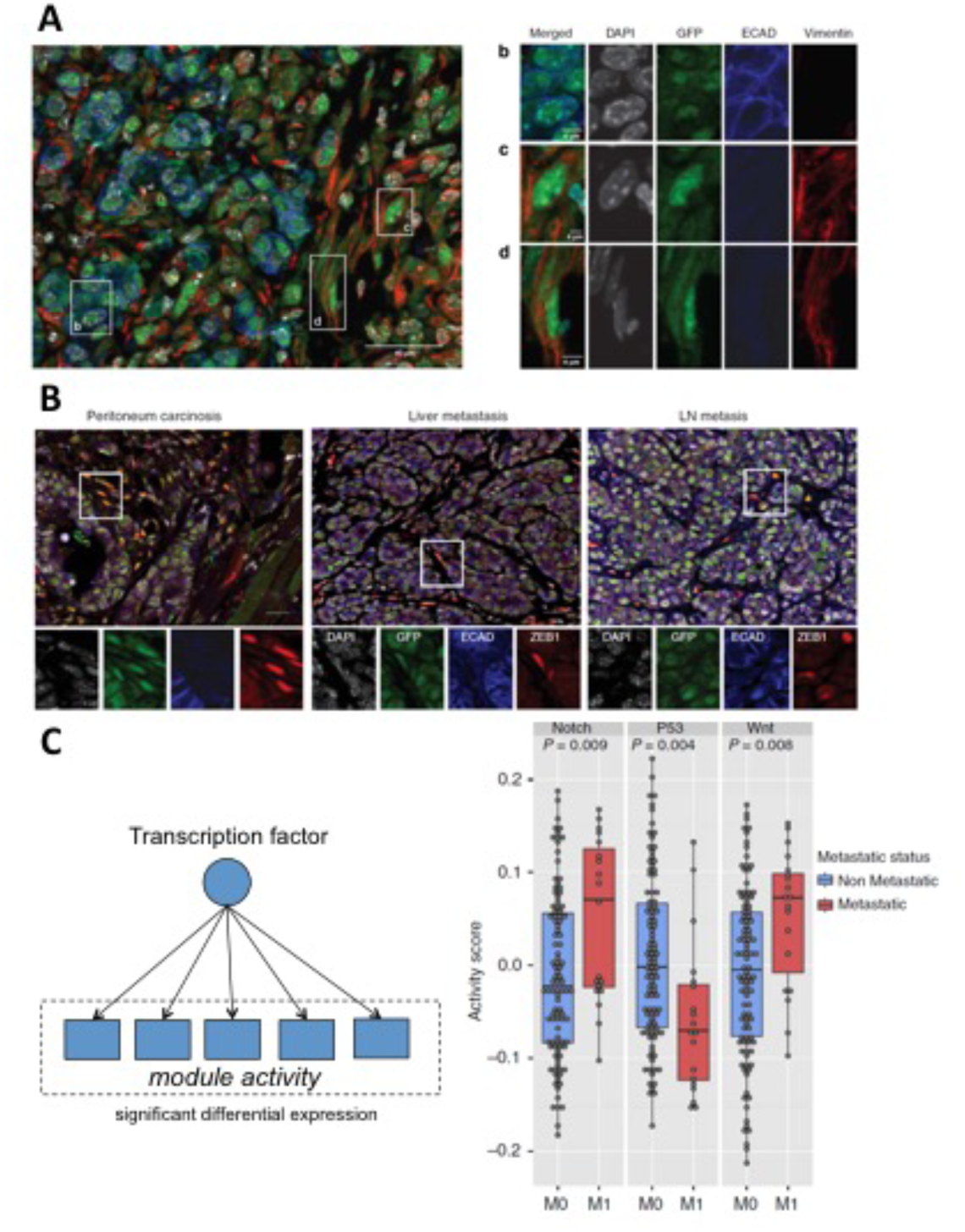
*p53 loss – Notch (NICD) overexpression double mutant results in invasive phenotype in colon cancer mice*. *Immunostaining for major EMT marker in (A) primary tumour and (B) metastases in distant organs; (C) Regulation of p53, Notch and Wnt pathways in invasive colon cancer in human (TCGA data). (Adapted from* [53]).

**Figure 11.**
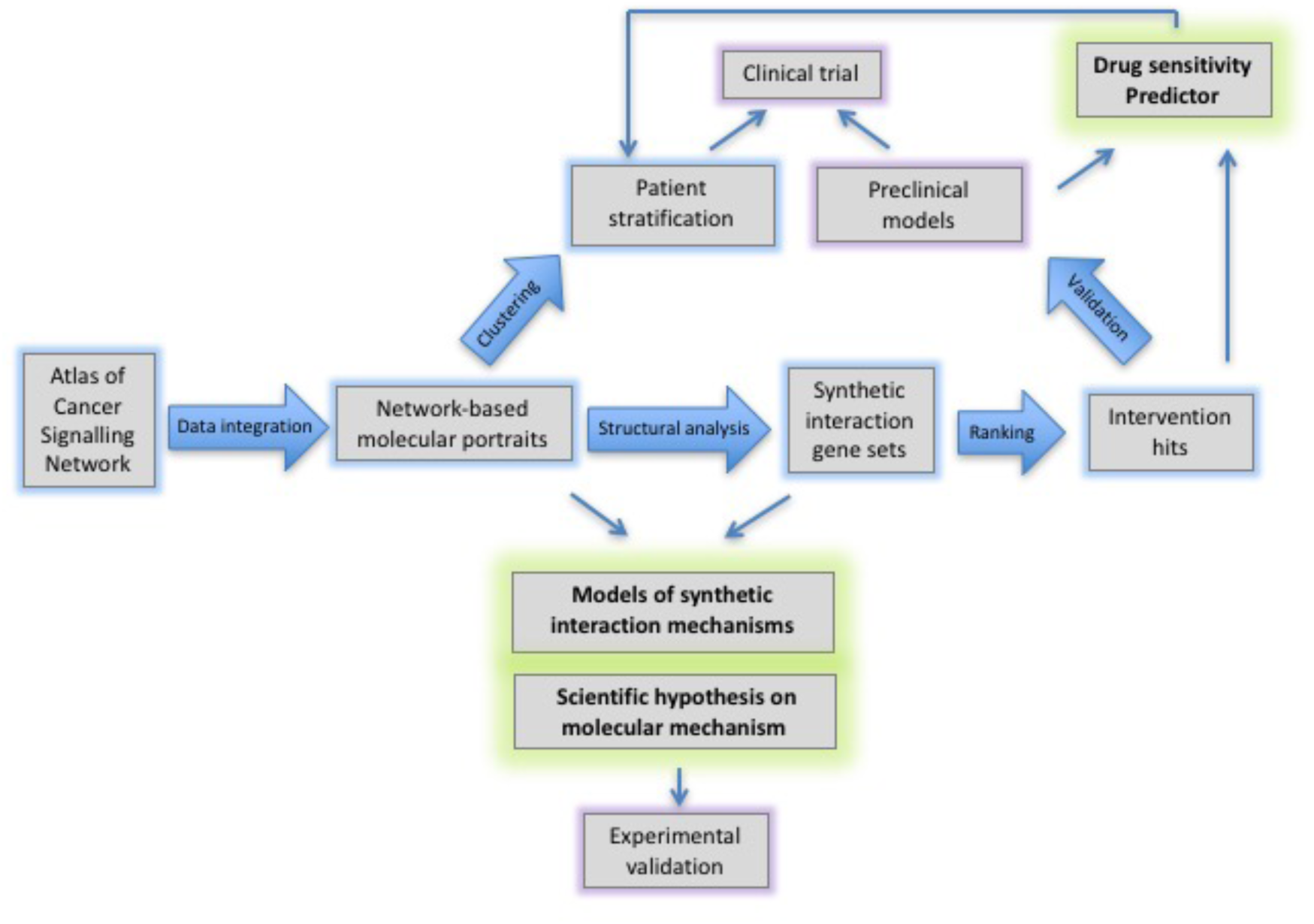
*Approaches for comprehensive cell signalling network maps applications together with high-throughput data*. *Approach can be useful for analysing perturbations in cellular processes and proposing synthetic combinations in research and in clinics: systematically represent molecular processes as comprehensive signalling networks, retrieve principles of signalling coordination and synthetic interactions, interpret omics data from disease in the context of signalling network, find signalling network fragilities in the disease, predict sensitivity to drug, suggest intervention hits*.

The prediction of synthetic interaction between Notch (NICD) and p53 demonstrated that there are alternative ways to reach permissive conditions to induce EMT, in addition to those already described in the literature. This idea was not intuitive and actually contradictory to the commonly accepted dogma in the colon cancer field. The study evokes an important message that gathering cell signalling mechanisms together may undercover un-expected interactions and lead to discovery of new mechanisms in regulation of cell phenotypes that may significantly affect the understanding of basic molecular processes implicated in cancer and change the therapeutic approaches. In addition, the comprehensive EMT signalling network is rich resource of information can be used in further studies. Finally, the new EMT mice is a relevant model mimicking the invasive human colon cancer and a system for therapeutic drugs discovery [54].

#### Finding susceptibility to papillary thyroid carcinoma development

Modules and maps of ACSN can serve as signatures of biological functions and can be used to find association between perturbations in particular molecular mechanisms to the risk of a specific cancer type development. In Lonjou et. al., the association of 141 SNPs located in 43 DNA repair genes from 10 DNA repair processes as it is depicted in the DNA repair map, was examined in 75 papillary thyroid carcinoma (PTC) cases and 254 controls. The study confirms that genetic variants in several genes operating in distinct DNA repair mechanisms are implicated in the development of PTC. In particular, a significant association of the intronic SNP rs2296675 of the *MGMT* gene from Direct Repair pathway with the risk of developing PTC was found [55]. Further investigation is undertaken, to decipher the molecular mechanisms controlled by the methyltransferase encoded by *MGMT* not only in Direct Repair pathway, but probably in other associated mechanisms, to understand how alteration of such functions may lead to the development of the most common type of thyroid cancer.

## Conclusions

This review is devoted to the use of signalling network modelling in cancer research, and exemplified with the multidisciplinary project Atlas of Cancer Signalling Network (ACSN). It described approaches addressing complexity of cancer by systematic representation of signalling implicated in the disease in the form of comprehensive signalling network maps and tools for network-based analysis, and visualisation and interpretation of cancer omics data. It summarized several studies on application of ACSN for finding synthetically interacting genes in cancer; predicting drug synergy; suggesting complex intervention sets and associating molecular mechanisms to cancer development susceptibility. These examples can serve as a basis for a more generalised approach. A fundamental question of signalling regulation in human disorders can be addressed by experimental and computational approaches, helping together to understand the principles of signalling rewiring [1]. It will have an impact on personalized intervention schemes, in particular those based on pharmacological combination. Organisation and formal representation of the knowledge of cell signalling followed by analysis of network features is a general paradigm that provides a more global view on molecular mechanisms and facilitates modelling of cell fates. It has a potential to explain mutant phenotypes, and reveal new molecular interactions. Considering the topology of signalling networks and studying network perturbations together with high-throughput data can guide toward an optimal therapeutic strategy in patients or in studied experimental systems, providing both new intervention points and relevant biomarkers.

There is ongoing effort to include ACSN content into aggregated pathway databases such as Pathway Commons [56] and WikiPathways [57] such that it can be used in many projects through standard interfaces as Cytoscape [58]. In addition, the approach adopted in ACSN projects has a wide potential, applicable for other complex diseases. This rationale has led to the creation of a collective research effort on different human disorders called Disease Maps project (http://disease-maps.org). This partnership aims at applying similar approaches as described in this review, that will lead to identification of emerging disease hallmarks. This will help to study diseases comorbidity, predict response to standard treatments and to suggest improved individual intervention schemes based on drug repositioning [6].

## Five key points summary

1. ACSN is a resource of cancer signalling knowledge, comprehensive map of molecular interactions in cancer based on the latest scientific literature.
2. NaviCell and NaviCom are interactive web-based environment for molecular maps navigation omics data integration and visualisation.
3. Interpretation of omics data from breast cancer cell lines with different sensitivity to targeted drugs in the context of signalling network to retrieve deregulated functional modules and to suggest synergy between drugs.
4. Application of signalling maps to find network fragilities in bladder cancer and to suggest intervention sets using OCSANA algorithm.
5. Prediction of non-intuitive combination of synthetically interacting genes in colon cancer using network analysis, formulating hypothesis on molecular mechanism and development of mice model.

## Acknowledgements

This work has been funded by the Agilent Thought Leader Award #3273 and ‘Projet Incitatif et Collaboratif Computational Systems Biology Approach for Cancer’ grant from Institut Curie. This work received support from APLIGOOGLE program provided by CNRS; the COLOSYS grant ANR-15-CMED-0001-04, provided by the Agence Nationale de la Recherche under the frame of ERACoSysMed-1, the ERA-Net for Systems Medicine in clinical research and medical practice and from INSERM Plan Cancer N° BIO2014-08 COMET grant under ITMO Cancer BioSys program.

